# Neuroimaging and plasma marker evidence for white matter macrostructure loss in Parkinson’s disease

**DOI:** 10.1101/2023.09.22.558937

**Authors:** Angeliki Zarkali, Naomi Hannaway, Peter McColgan, Amanda J Heslegrave, Elena Veleva, Rhiannon Laban, Henrik Zetterberg, Andrew J Lees, Nick C. Fox, Rimona S Weil

## Abstract

Parkinson’s disease (PD) is the second commonest neurodegenerative disorder and over half of patients progress to postural instability, dementia or death within 10 years of diagnosis.

However, onset and rate of progression to poor outcomes is highly variable, underpinned by heterogeneity in the underlying pathological process. Improved biomarkers of poor outcomes would be helpful for targeted treatment, but most studies to-date have been limited to a single modality or the assessment of patients with established cognitive impairment. Here, we use multimodal neuroimaging and plasma biomarkers in 98 patients with PD and 28 age-matched controls followed-up over 3 years, including: gray matter (cortical thickness), white matter (macrostructure: fibre-cross section and microstructure: fibre density) at whole-brain and tract level, structural and functional connectivity and plasma levels of neurofilament light chain (NFL) and phosphorylated tau (p-tau) 181. We show extensive reductions in fibre cross-section and structural connectivity in PD with poor outcomes, with preserved gray matter and functional connectivity. NFL, but not p-tau181 levels was increased in PD with poor outcomes and correlated with white matter loss. These findings suggest that imaging sensitive to white matter macrostructure and plasma NFL may be useful biomarkers of poor outcomes in PD. As new targeted treatments are emerging, these biomarkers show important potential to aid patient selection for treatments and improve stratification to clinical trials.

## Introduction

Parkinson’s disease (PD) is the second commonest neurodegenerative condition^1^. As well as the well-described motor symptoms of rest tremor, rigidity and bradykinesia, around half of all patients will develop dementia within 10 years’ of diagnosis^2^. Other poor outcomes include frailty and falls due to postural instability; and PD dementia has higher societal and economic burden than other dementias^3,4^. However the timing and rate of clinical deterioration varies greatly^2,5^ as does the underlying brain pathology, with varying degrees and locations of alpha-synuclein, and the extent and severity of beta-amyloid and tau pathological accumulations^6^. Although factors such as older age, male sex, baseline cognition, particularly visuoperceptual function,^7^ are associated with poor clinical outcomes, and clinical algorithms to predict risk are being developed^8,9^, underlying changes in brain structure and function in these patients at greater risk remain unclear.

Most studies examining neuroimaging changes in association with poor outcomes in PD have been limited to a single modality, mainly focusing on measures of gray matter, and the results have been inconsistent^10^.Both frontal^11^ and temporoparietal cortical thickness changes^12^ have been reported in patients with PD who subsequently develop dementia, with several other areas implicated^13,14^. However, gray matter changes reflect neuronal loss^15^, which only occurs later in the disease process^16^ making it a less reliable marker of severity in Parkinson’s disease.

Instead, evidence from animal models has shown that white matter degeneration precedes neuronal loss in PD^17,18^, suggesting that in-vivo markers of white matter integrity, rather than gray matter, might be useful imaging biomarkers of severity in PD. Chung et al, recently showed white matter alterations in PD patients with mild cognitive impairment (PD-MCI) who progressed to develop dementia. Several tracts, were implicated, including the arcuate fasciculus bilaterally and the left cingulum^19^. However, that study evaluated patients with established cognitive impairment, and used only diffusion tensor imaging analysis which cannot accurately model fibers with divergent orientations (which make up the majority of fibre tracts in the brain)^20^.

Higher tensor diffusion models have been recently developed. These provide more accurate measures of crossing fibers, allowing better estimation of white matter *in vivo*. Fixel-based analysis is one such model, and was applied by Rau et al. to reveal reduction in fibre cross-section within the anterior body of the corpus callosum in PD patients with more severe disease^21^. We recently used fixel-based analysis to show widespread changes in PD patients with poor visuoperceptual function, who are at higher risk of dementia^22^. Changes at the whole network level may also be a more sensitive marker of poor outcome in PD: reduction in structural connectivity is seen in patients with PD-MCI who subsequently progress to dementia^19^, whilst long-range interhemispheric connections are more affected in PD patients at higher risk of dementia^23^.

Complementary to imaging biomarkers, biomarkers in the plasma and cerebrospinal fluid are becoming more readily available and provide insights into disease severity and underlying processes. Neurofilament light chain (NFL) reflects axonal damage and shows higher CSF concentrations relating to white matter lesions in conditions including multiple sclerosis and Alzheimer’s^24,25^. CSF NFL is higher in patients with PD with established cognitive impairment^26^ and plasma NFL is increased in PD patients who later developed PD-MCI or PD dementia^27,28^. In contrast to NFL, which relates to axonal damage, disease-relevant pathological accumulations can now be detected at very low concentrations in the plasma. Plasma levels of phosphorylated tau at threonine 181 (p-tau181) is now established in Alzheimer’s and other dementias as a marker of tau as well as β-amyloid pathology^29^ and is of relevance in PD, as β-amyloid plaques and tau deposition are seen in over three quarters of patients with PD dementia at post-mortem^30^. In PD patients who progress to dementia, p-tau181 levels were not increased, although in the related condition, dementia with Lewy bodies, higher levels of plasma p-tau181 correlated with greater degree of cognitive decline^28,31^. The relationship between plasma markers such as NFL and p-tau181, with neuroimaging changes in patients with PD, and how these relate to poor clinical outcomes is not yet clear.

Establishing relative changes in imaging and plasma measures in patients with PD who go on to develop poor clinical outcomes can provide key insights into the sequence of underlying pathological changes in these patients. Specifically, higher tensor models of diffusion imaging, and plasma NFL can shed light on the role of axonal damage; whereas plasma p-tau181 provides information about brain levels of tau and beta-amyloid. At the same time, biomarkers predictive of poor outcomes in PD can improve efficiency and stratification for clinical trials, as well as more targeted treatment, as new pathology-specific interventions emerge.

Here, we examined neuroimaging and plasma biomarkers in 98 patients with PD followed up over 3 years (*Figure 1*). Using multimodal neuroimaging at baseline, we assessed: a) gray matter changes using cortical thickness, b) white matter microstructural and macrostructural changes at whole-brain and tract level using fixel-based analysis, a technique able to reliably model crossing fibers^32^ and c) changes in functional and structural connectivity at whole-network level in patients with PD who have poor outcomes during follow-up compared to those with PD and good outcomes. In addition, we assessed differences in the concentration of two plasma biomarkers (NFL and plasma p-tau181) taken during follow-up sessions in patients with PD and poor outcomes, compared to those with good outcomes; and we investigate the relationship between plasma and imaging biomarkers.

**Figure 1.**
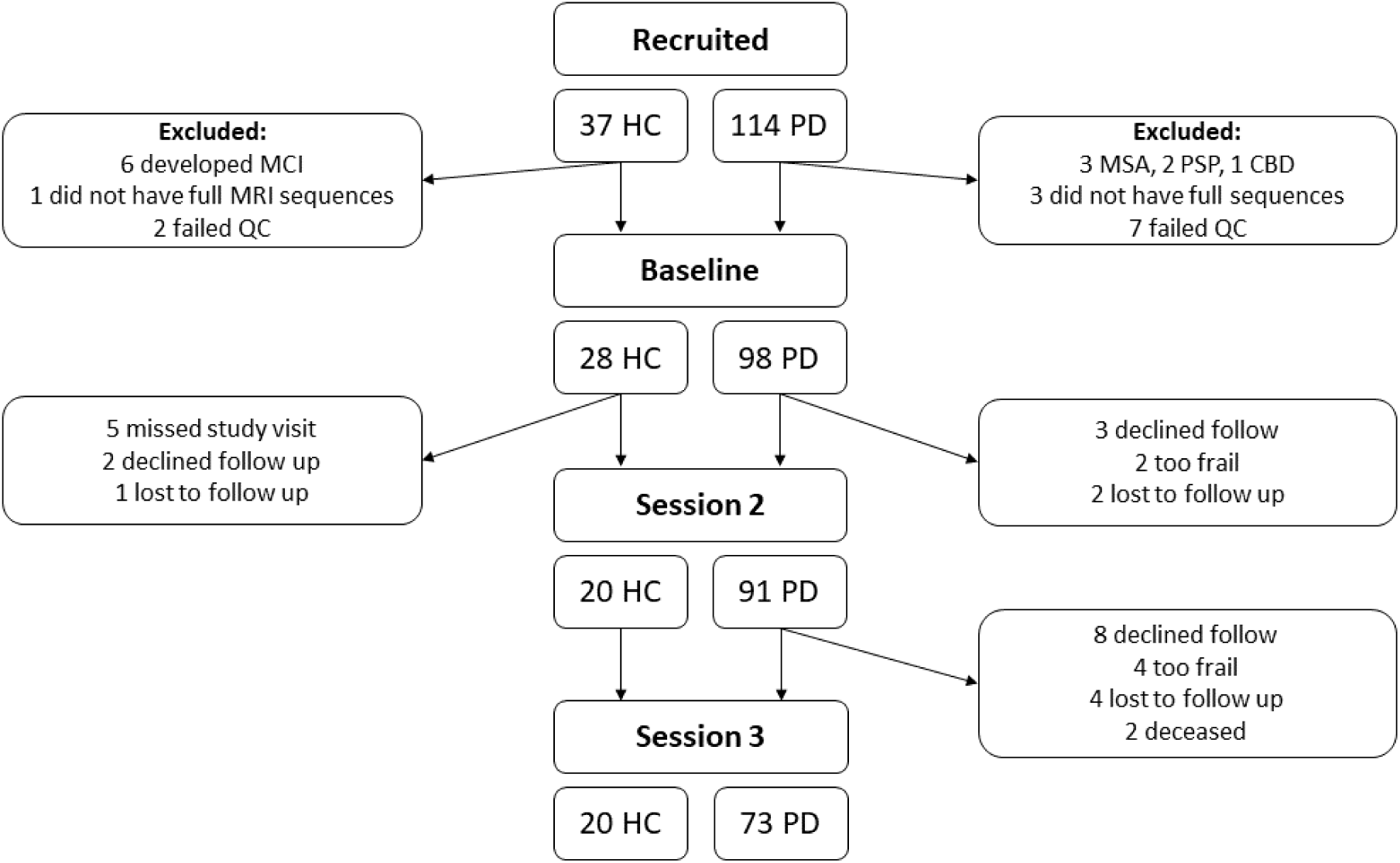
Overview of the recruited participants. A total of 114 patients with Parkinson’s (PD) and 37 healthy controls (HC) were recruited. After exclusion of PD patients with a change in diagnosis to atypical parkinsonism, controls who developed mild cognitive impairment (MCI) and those who did not complete the full scanning sequence or failed quality control (QC), a total of 98 patients with PD and 28 HC were included. All participants underwent MRI imaging at baseline (including T1-weighted imaging, diffusion weighted imaging and resting state functional MRI). At Session 2, participants had bloods taken for plasma biomarkers (neurofilament light and phosphorylated tau). At all study sessions participants underwent clinical and cognitive assessments. CBD: corticobasal degeneration, MSA: multiple system atrophy, PSP: progressive supranuclear palsy,

## Results

A total of 98 people with PD and 28 healthy controls were included at baseline. 8 PD participants could not attend further follow-up due to frailty (2 at 18 months, 4 at 3 years) or death (2 participants at 3 years). A further 11 PD and 2 control participants declined further follow up and 6 PD and 1 control participants were lost to follow-up (*Figure 1*). PD participants were defined as PD poor outcome if death, frailty, dementia or mild cognitive impairment (MCI) developed at any point during follow-up.

Results of demographics and baseline clinical assessments are seen in *Table 1*. Importantly, the groups did not significantly differ in terms of MRI image quality metrics. PD with poor outcome were older at baseline (mean ± std: 68.5 ± 8.5 years in PD poor outcome vs 62.4 ± 7.0 in PD good outcome), with higher male predominance (74.2% in PD poor outcome vs 43.3% in PD good outcomes), higher depression scores (5.2 ± 3.3 in PD poor outcome vs 3.4 ± 2.7 in PD good outcome), albeit below the clinical threshold =<8. They also performed worse than PD good outcome and controls in visual assessments including colour (0.016) and contrast sensitivity (p=0.006) and in cognitive testing including MMSE (p-0.004), MOCA (p<0.001), Stroop Colour (p=0.002), Stroop Interference (p=0.001), Verbal fluency category (p=0.003), Delayed Logical memory (p<0.001), Judgement of line orientation (p=0.042) and Hooper task (p<0.001). Finally, PD poor outcomes had higher baseline UPDRS total (p=0.007) and UPDRS motor scores (p=0.042) than PD good outcomes (*Table 1*). Longitudinal change in cognition and disease-specific measures are presented in *Table 2*.

**Table 1.**
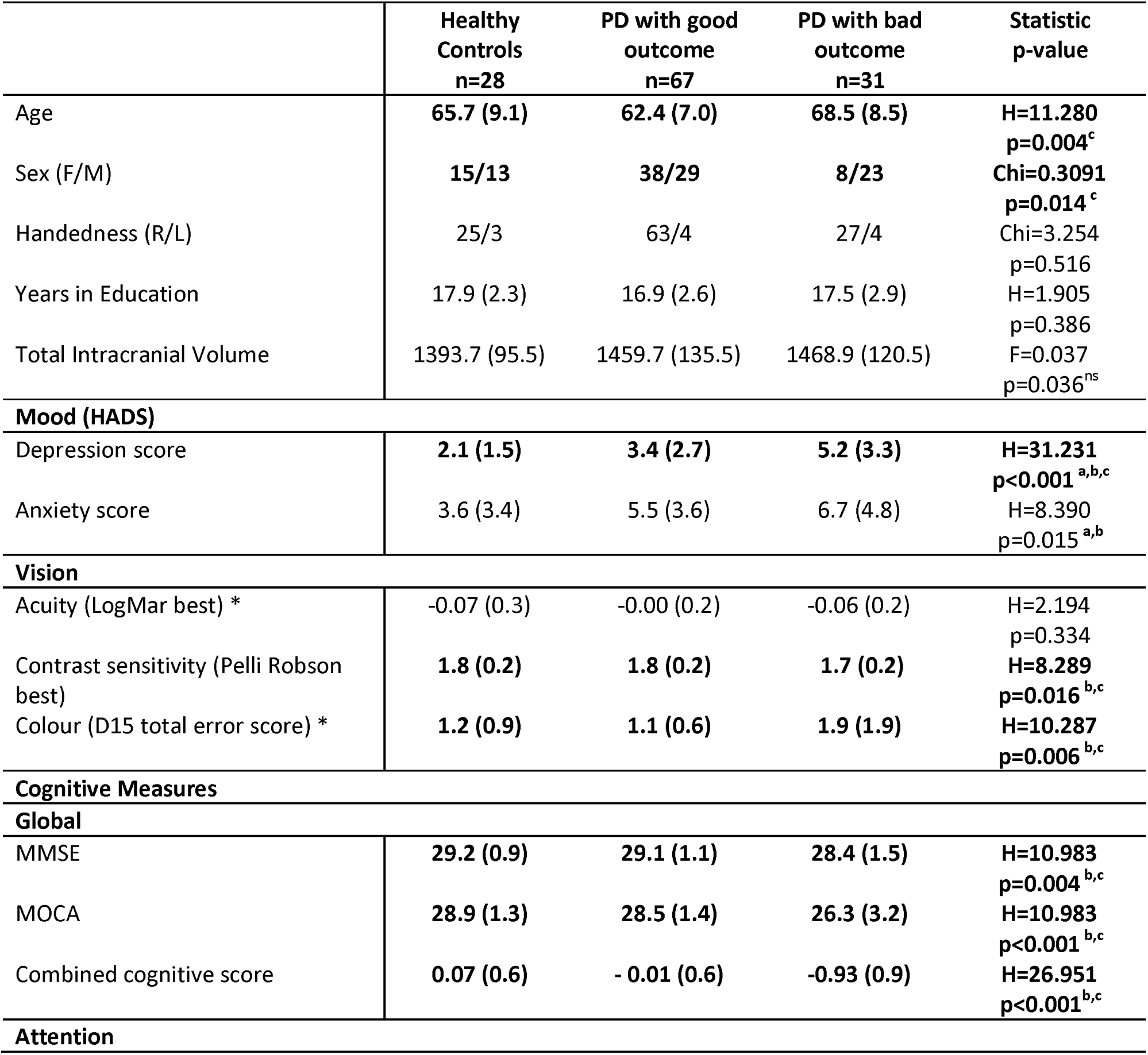
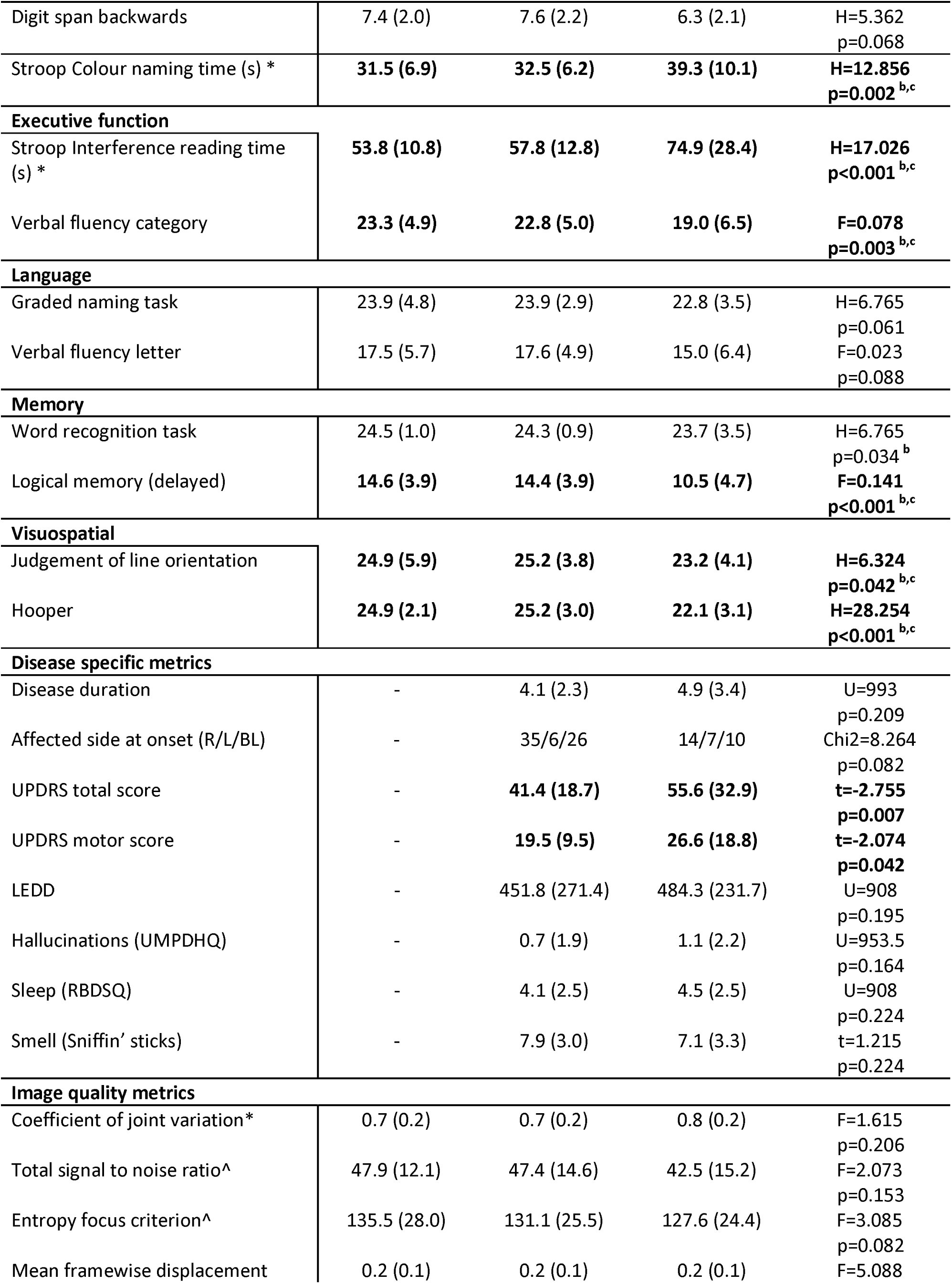
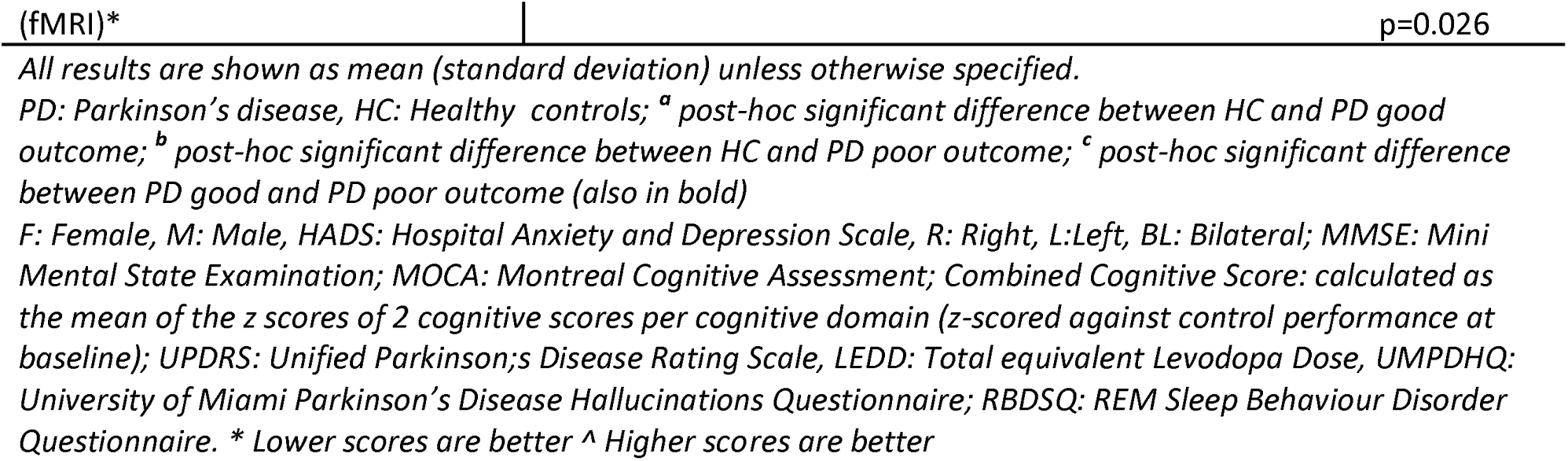
Baseline demographics and clinical assessments.

**Table 2.**
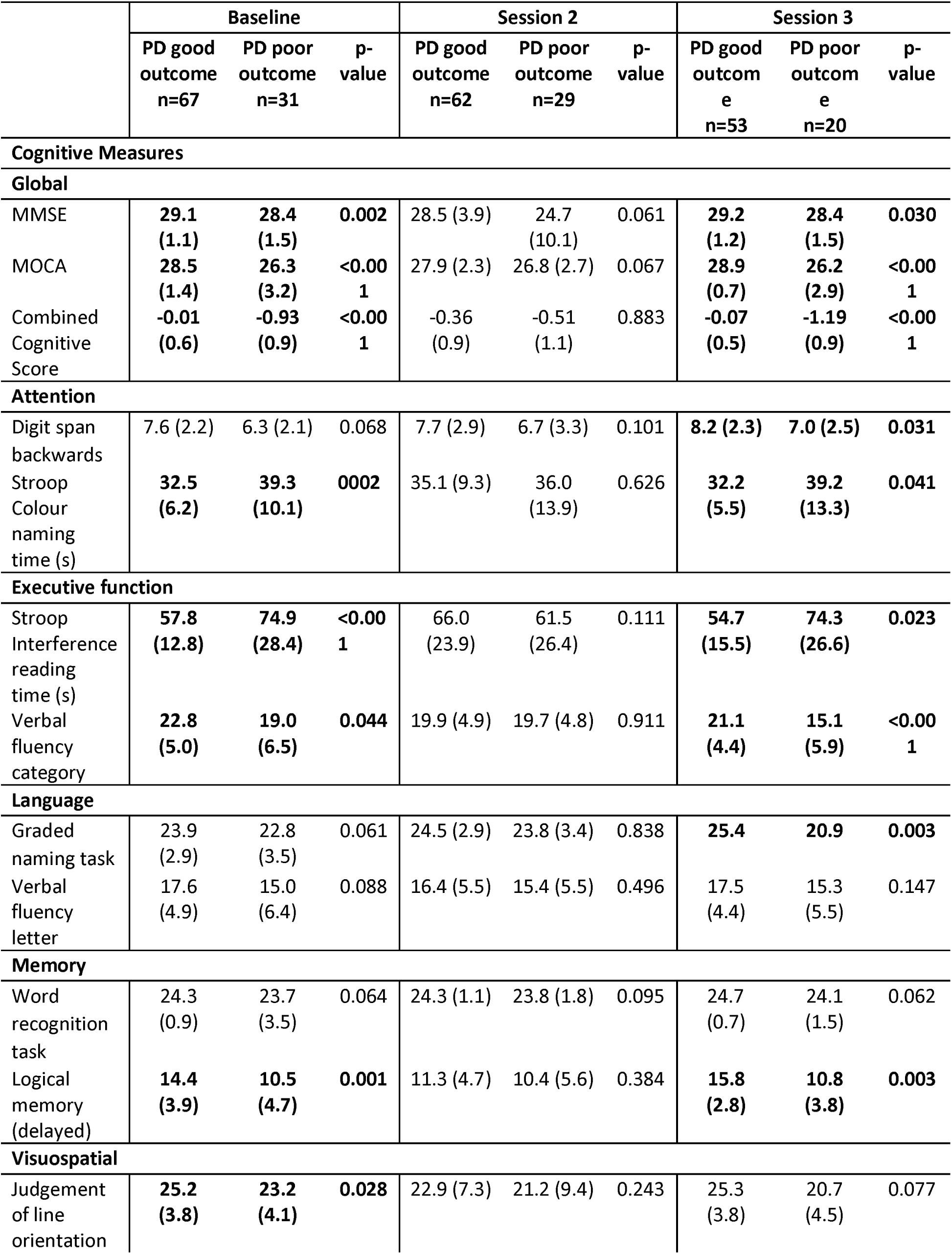
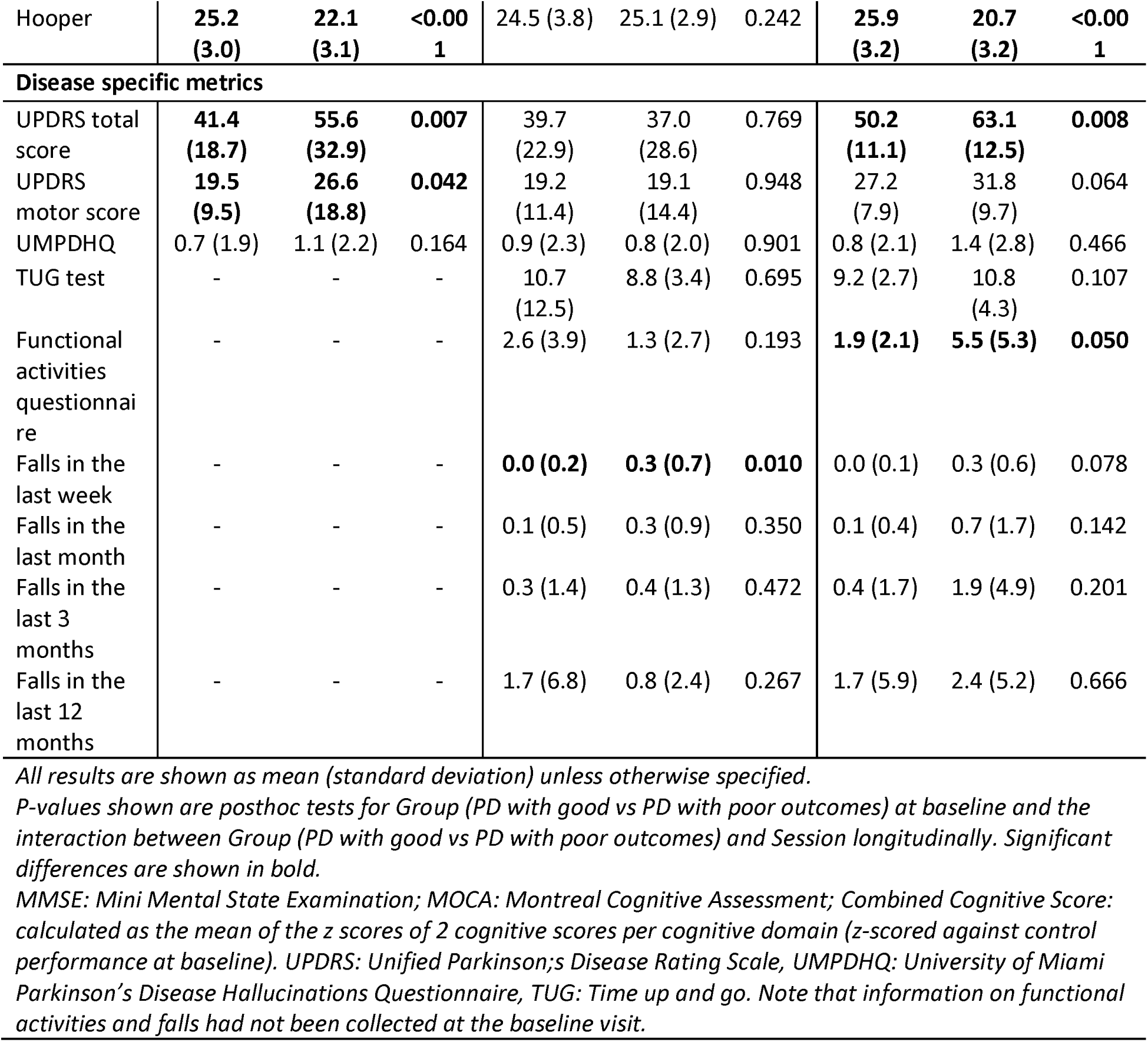
Longitudinal change in cognitive and disease specific metrics in patients with Parkinson’s disease (PD).

### Extensive macrostructural white matter changes at baseline in PD with subsequent poor outcome, in the absence of gray matter changes

No significant differences were seen in cortical thickness at whole brain level (adjusting for age sex and total intracranial volume) between PD patients with poor vs PD patients with good clinical outcomes at baseline.

We then assessed, white matter micro-structure (fibre density FD), macro-structure (fibre cross-section FC) and overall white matter integrity (combined fibre density and cross-section FDC) at baseline at whole-white-matter level in PD patients with poor vs PD patients with good outcomes, adjusting for age, sex and total intracranial volume (for FC and FDC). We found extensive macrostructural changes at baseline with up to 19% reductions in PD poor outcome compared to those with good outcomes in several tracts: bilateral anterior thalamic radiations, optic radiations, inferior fronto-occipital fasciculi, cingulum, thalamo-prefrontal and thalamo-parietal tracts, left corticospinal tract, left middle longitudinal fasciculus, left superior thalamic radiation, left thalamo-parietal tract, left, corpus callosum and middle cerebellar peduncle (*Figure 2A*). In contrast, white matter microstructure (FD) and FDC did not significantly differ amongst PD patients.

**Figure 2.**
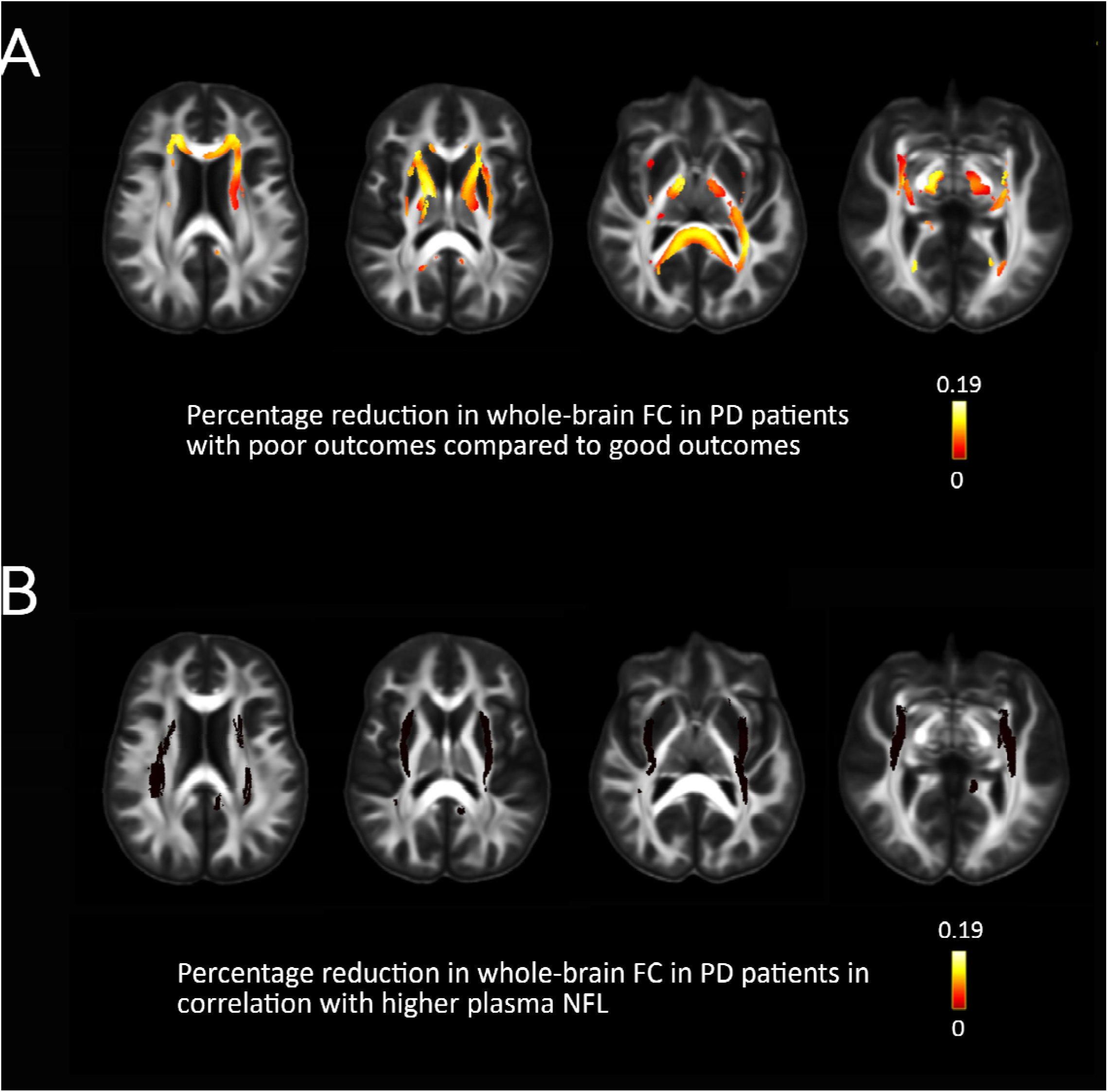
White matter macrostructural changes in patients with Parkinson’s disease (PD) and poor outcomes. A. White matter macrostructural changes (reduction in fibre cross-section (FC)) in PD patients with poor outcomes compared to PD patients with good outcomes at baseline whole-white-matter analysis with age, sex and total intracranial volume as nuisance covariates. B. Reductions in FC in PD patients in relation to higher plasma NFL values. Effect sizes are shown as percentages, presented as streamlines, only family-wise-error (FWE) corrected results are displayed (FWE-corrected p<0.05).

We then assessed mean fibre cross section across 52 white matter tracts using TractSeg between patients with PD with good vs with poor outcomes, adjusting for age, sex and total intracranial volume. We found reduced mean FC in PD with poor outcomes within the left arcuate fasciculus, left anterior thalamic radiation, right medial longitudinal fasciculus, left optic radiation, and left thalamo-prefrontal tract. Reduced mean FC was also seen within the corpus callosum, driven by reductions within the genu, posterior midbody and splenium, whilst increased mean FC was seen within the rostral body of the corpus callosum and the left thalamo-occipital tract (*Figure 3A*). Results for all 52 tracts included in the analyses are presented in *Supplementary Figure S2*. All of the tracts that were significantly different in PD with poor outcomes were significantly correlated with combined cognitive scores at baseline (FDR-corrected, adjusting for age, sex and total intracranial volume) and all tracts that showed reduced mean FC in PD with poor outcomes were significant correlated with combined cognitive scores at last follow up (FDR-corrected, adjusting for age, sex, total intracranial volume and time-to-follow-up) (*Table 3*). Mean FC within the genu of the corpus callosum and the left anterior thalamic radiation were correlated with longitudinal change in combined cognitive scores, but neither survived correction for multiple comparisons (*Table 3*).

**Figure 3.**
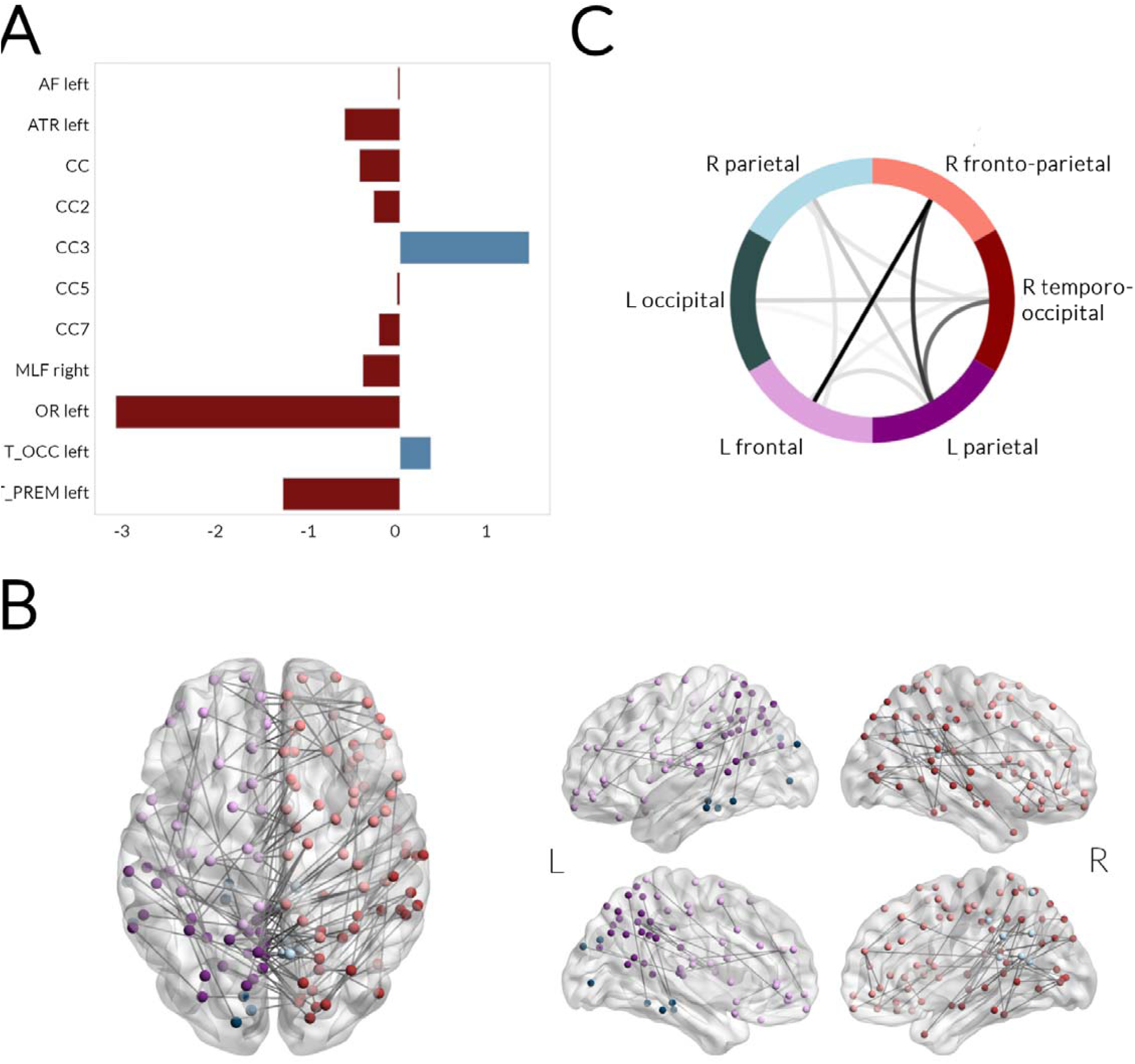
Changes in tract macrostructure and structural connectivity in patients with Parkinson’s disease (PD) and poor outcomes. A. Tract of interest analysis. Mean fibre cross-section (FC) along 52 white matter tracts, segmented using TractSeg, were compared between PD with poor outcomes vs PD with poor outcomes at baseline, correcting for age, sex and total intracranial volume, false discovery rate (FDR) corrected for multiple comparisons. PD with poor outcomes showed reduction in mean fibre cross-section (FC) in the left arcuate fasciculus (AF), left anterior thalamic radiation (ATR), corpus callosum (CC), specifically the genu (CC2), posterior midbody (CC5) and splenium (CC7), the right medial longitudinal fasciculus (MLF), left optic radiation (OR) and left thalamo-prefrontal tract (T_PREM). Increased mean tract FC was seen in PD with poor outcomes in the rostral body of the corpus callosum (CC3) and the left thalamo-occipital tract (T_OCC). All results presented are FDR-corrected p<0.05, presented as percentage change from PD with good outcomes. B. Structural connectivity changes. Network based statistical analysis revealed a network of reduced connectivity strength in PD with poor outcomes (FDR-corrected p<0.05, t=3.0, 5000 permutations, correcting for age and sex), which comprised 215 edges and 105 nodes across 6 modules. The subnetwork was visualized using BrainNetViewer with different colors for each module. C. Between modules connectivity changes. The network of reduced structural connectivity in PD with poor outcomes comprised of 6 modules: R parietal, R fronto-parietal, R temporo-occipital, L parietal, L frontal and L occipital. The sum number of connections between modules showing reduced connectivity strength is visualised with darker colour. Connection within R fronto-parietal and L frontal, R frontoparietal and L parietal and R temporo-occipital and L parietal modules were most affected in PD with poor outcomes. L: left, R: right

**Table 3.**
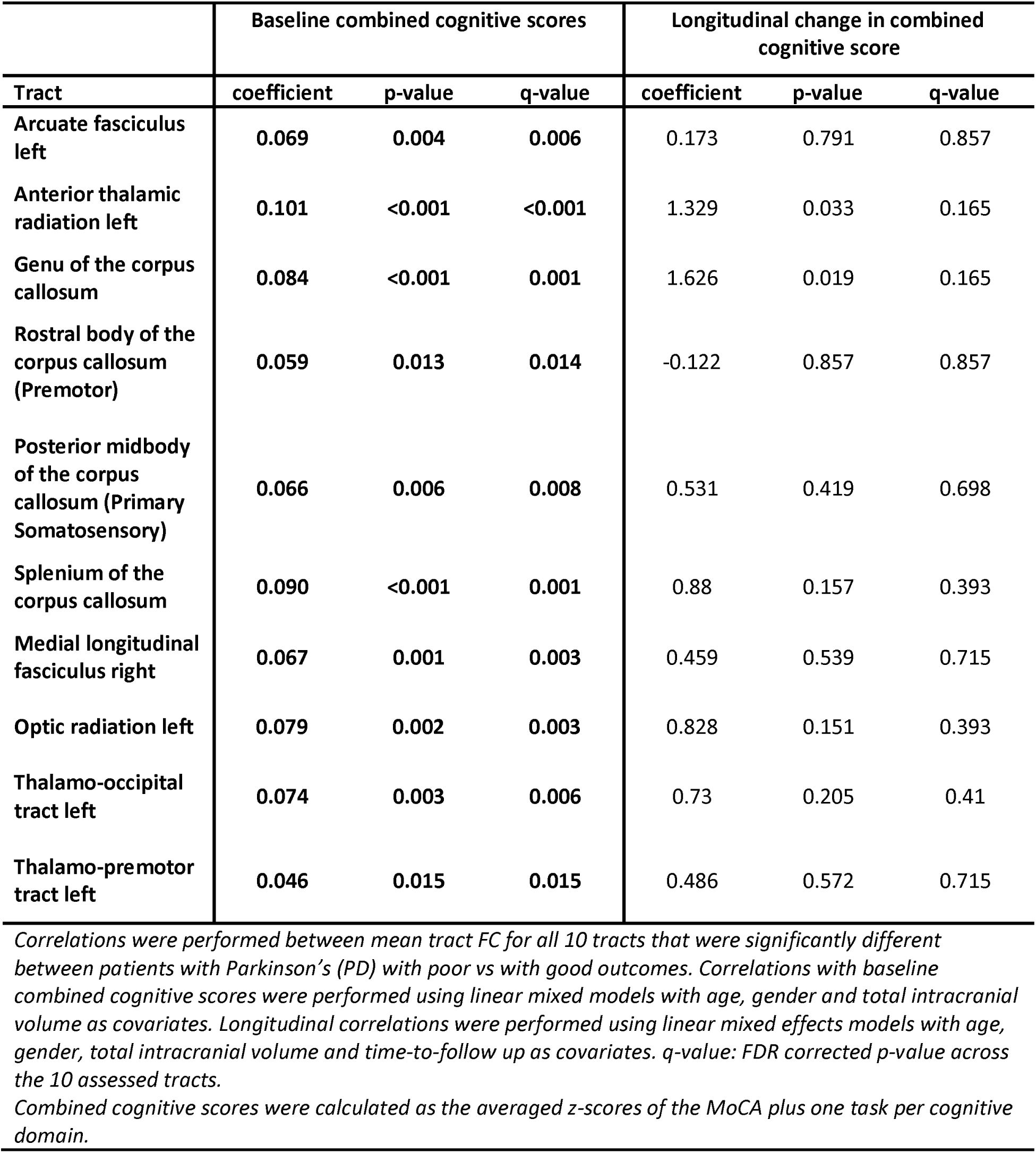
Correlation between tracts of interest showing reduced mean fibre cross section (FC) in PD patients with poor outcomes and cognition, at baseline and longitudinally.

| Tract | Baseline combined cognitive scores |  |  | Longitudinal change in combined cognitive score |  |  |
| --- | --- | --- | --- | --- | --- | --- |
|  | coefficient | p-value | q-value | coefficient | p-value | q-value |
| Arcuate fasciculus left | 0.069 | 0.004 | 0.006 | 0.173 | 0.791 | 0.857 |
| Anterior thalamic radiation left | 0.101 | <0.001 | <0.001 | 1.329 | 0.033 | 0.165 |
| Genu of the corpus callosum | 0.084 | <0.001 | 0.001 | 1.626 | 0.019 | 0.165 |
| Rostral body of the corpus callosum (Premotor) | 0.059 | 0.013 | 0.014 | -0.122 | 0.857 | 0.857 |
| Posterior midbody of the corpus callosum (Primary Somatosensory) | 0.066 | 0.006 | 0.008 | 0.531 | 0.419 | 0.698 |
| Splenium of the corpus callosum | 0.090 | <0.001 | 0.001 | 0.88 | 0.157 | 0.393 |
| Medial longitudinal fasciculus right | 0.067 | 0.001 | 0.003 | 0.459 | 0.539 | 0.715 |
| Optic radiation left | 0.079 | 0.002 | 0.003 | 0.828 | 0.151 | 0.393 |
| Thalamo-occipital tract left | 0.074 | 0.003 | 0.006 | 0.73 | 0.205 | 0.41 |
| Thalamo-premotor tract left | 0.046 | 0.015 | 0.015 | 0.486 | 0.572 | 0.715 |

*Correlations were performed between mean tract FC for all 10 tracts that were significantly different between patients with Parkinson's (PD) with poor vs with good outcomes. Correlations with baseline combined cognitive scores were performed using linear mixed models with age, gender and total intracranial volume as covariates. Longitudinal correlations were performed using linear mixed effects models with age, gender, total intracranial volume and time-to-follow up as covariates. q-value: FDR corrected p-value across the 10 assessed tracts.*
*Combined cognitive scores were calculated as the averaged z-scores of the MoCA plus one task per cognitive domain.*

### Plasma NFL but not p-tau181 levels are increased in PD patients with poor outcomes and is correlated with white matter macrostructure

PD patients with poor outcomes had higher levels of plasma NFL adjusting for age and sex compared to PD patients with good outcomes (β=4.378, p=0.016) but levels of plasma p-tau181 were not significantly different between groups, adjusting for age, sex and batch (β=0.461, p=0.106) (*Figure 4A,B*). Mean FC of the areas showing macrostructural changes in PD with poor outcomes was significantly negatively correlated with plasma NFL concentration (rho=-0.436, p<0.001) but not p-tau181 levels (rho=-0.153, p=0.157) (*Figure 4C,D*). In addition, within patients with PD, higher plasma NFL concentration was associated with lower FC (up to 1% reductions) at whole brain analysis within bilateral inferior fronto-occipital fasciculi and optic radiations (*Figure 2B*).

**Figure 4.**
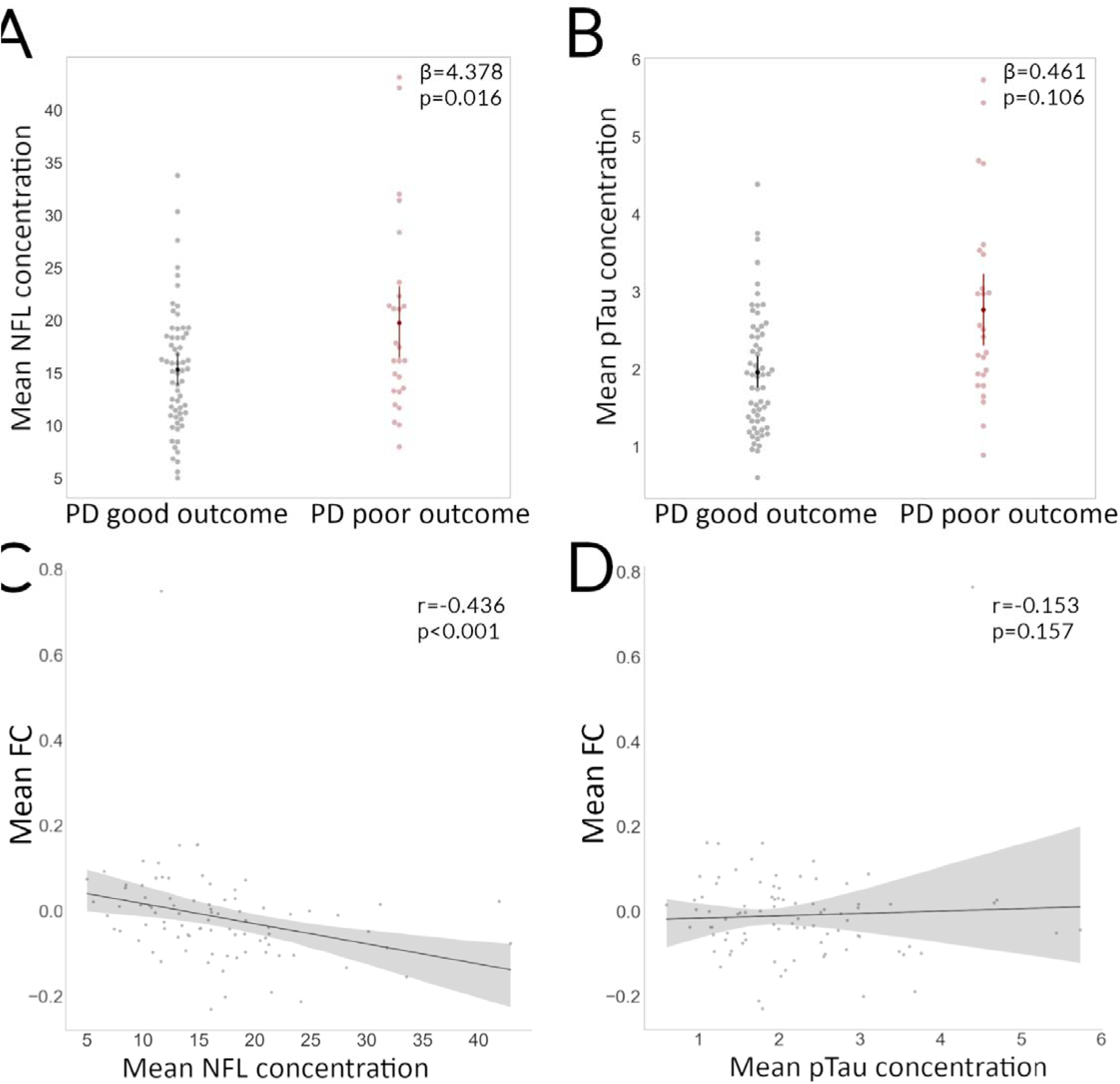
Changes in plasma biomarkers in patients with Parkinson’s disease (PD) and poor outcomes. A. PD patients with poor outcomes show increased plasma neurofilament light chain (NFL) concentration, adjusting for age and sex, compared to PD with good outcomes. B. PD with poor outcomes show increased plasma concentration of phosphorylated tau at threonine 181(p-tau181), adjusting for age and sex, compared to PD with good outcomes. C. Mean fibre cross-section (FC) in areas showing significant whole-white-matter reductions in PD with poor outcomes was significantly correlated with plasma NFL concentration within patients with PD. D. Mean fibre cross-section (FC) in areas showing significant whole-white-matter reductions in PD with poor outcomes was not significantly correlated with plasma p-tau181 concentration within patients with PD.

### Structural but not functional connectivity is reduced in PD patients with poor outcomes

Finally, we assessed baseline changes at network level in PD patients who had poor outcomes at follow-up, using both functional connectivity from rsfMRI and structural connectivity derived from diffusion weighted imaging. Functional and structural connectomes were compared between PD patients with poor vs good outcomes using network-based statistics, with age and sex included as nuisance covariates (5000 permutations, t=3.0, FDR-corrected value <0.05). Whist there were no group differences in functional connectivity, we found significant changes in structural connectivity, with reduced connectivity within a network involving 105 nodes and 235 edges and consisting of 6 modules (*Figure 3B*). Connections between the right fronto-parietal and left frontal, right fronto-parietal and left parietal and right temporo-occipital and left parietal modules were most affected in PD with poor outcomes (*Figure 3C*).

## Discussion

In this study we examine the changes in neuroimaging and plasma biomarkers in patients with PD who go on to develop poor clinical outcomes. We show that extensive macrostructural white matter changes can be detected in patients with PD who will progress to poor outcomes, with up to 19% reduction in fibre cross-section across multiple tracts, and widespread reductions in structural connectivity; and increased levels of plasma biomarkers, particularly NFL. In contrast, cortical thickness, white matter microstructure and functional connectivity are not significantly different amongst patients with PD with poor outcomes compared to those who have good outcomes.

This finding provides evidence that axonal changes are likely to be important in the pathophysiology of poor outcomes in PD and that white matter imaging is a more valuable and sensitive marker of poor outcomes than gray matter imaging. Specifically, we show macrostructural changes (reductions in FC) but preserved microstructure (FD). In a recent study of Alzheimer’s disease, macrostructure was related to pathological beta-amyloid and tau accumulation on PET imaging, whilst microstructural measures such as FD were correlated with the presence of white matter hyperintensities. Reduced FC has been previously shown in PD compared to healthy controls^21^ and amongst those with poor visual performance who are at a higher risk for dementia^22^. Although we did not have imaging measures sensitive to small vessel disease in our cohort, our current study provides further evidence of specific macrostructural alterations with preserved microstructure in PD. Future work, with appropriate imaging sequences, could specifically examine the link between small vessel disease and white matter microstructure in PD.

Similarly, despite the significant, extensive changes in structural connectivity, functional connectivity could not separate PD patients with poor vs good outcomes at baseline. This could reflect compensatory changes in functional connectivity. In the aging brain, despite an overall reduction in streamlines^34^ and reduced myelin integrity^35^ the functional connectome undergoes extensive reorganisation across a posterior-anterior gradient^36^ with both increases and reductions in functional connectivity^36,37^ compensating for the change in structural connectivity and initially preserving cognitive function^38^. Changes in temporal dynamics rather than static functional connectivity are more sensitive to structural changes accompanying aging^39^ and have been shown to be altered in PD with mild cognitive impairment and dementia^40^. Longitudinal assessment of functional connectivity incorporating approaches examining temporal dynamics may be more sensitive as an early biomarker of poor outcomes in PD.

Our network analysis revealed extensive structural connectivity changes in PD patients with poor outcomes. Interhemispheric connectivity between frontal, parietal and right occipital regions was most affected. Prior studies have shown structural connectivity changes in patients with established PD-MCI^41,42^ and more recently, patients with PD-MCI who later went on to have PD dementia were found to show further reductions in both frontal and occipital regions compared with PD patients who did not become demented^19^. Our findings also confirm both frontal and posterior changes in structural intra-but mostly interhemispheric connectivity, and reveal that these changes are seen in susceptible individuals even before the onset of PD dementia.

In addition to neuroimaging measures, we assessed plasma NFL and p-tau181 levels. NFL but not p-tau181 levels were increased in PD patients with poor outcomes, although these were found at follow up, rather than at baseline, when several patients had already developed poor outcomes. NFL was additionally correlated with white matter macrostructural changes with reduced FC within the left inferior fronto-occipital fasciculus at whole-brain analysis in association with higher plasma NFL concentration. Higher NFL concentration in the CSF has been correlated with shorter survival and worse motor symptoms in PD^43^ and higher plasma NFL values have been shown in patients with PD and established cognitive impairment^27,28,44^, correlating with rates of cognitive decline^28^. Our finding of a relationship between plasma NFL and loss of white matter microstructural integrity is consistent with NFL being a measure of axonal damage^24,25^.

In contrast, similar to other groups, we did not find a relationship between plasma p-tau181 and cognition in PD^28,44^. It is notable that in patients with DLB or at more advanced stages of PD dementia, higher p-tau181 was found to correlate with more rapid cognitive decline ^45^. This could suggest that axonal changes are earlier events in the progression from PD to PD dementia, with pathological accumulation of brain beta amyloid and tau occurring at later stage.

Plasma NFL may have a role as a predictive biomarker for poor outcomes in PD, and may be used together with imaging measures of structural white matter integrity. An alternative explanation for our lack of correlation between p-tau181 and PD cognition might be that patients vary in the extent of beta-amyloid and tau accumulation in the brain. Instead, p-tau181 may have a more useful role in identifying which patients have higher levels of these proteins and be future candidates for specific efficacious anti-amyloid (or anti-tau) therapies.

Finally, our study also highlights demographic risk factors of poor outcomes, including older age, male gender and poorer visuoperceptual function^7^. Specifically, older age at onset has been highlighted by several epidemiological and pathological studies as a risk factor for poorer clinical outcome and more rapid rate of progression in PD^46–49^. Beta-amyloid and tau accumulates with aging, even in cognitively intact individuals^50,51^, and so does cerebrovascular disease^52^. Increased inflammation^53^, impaired autophagy and protein clearance^54^, mitochondrial dysfunction^55^ and impaired DNA repair^56^ also accompany aging. How these age-related changes interplay in combination with alpha-synuclein pathology in PD still remains unclear.

### Limitations and Future Directions

Our study had some limitations. We followed patients for 3 years, and classified patients as having poor outcomes by the last follow-up session. Inevitably, some patients classified as not having poor outcomes will go on to have poor outcomes with longer follow up. Our MRI sequences did not include T2 or FLAIR sequences, which would be required to quantify concurrent small vessel disease, which is likely to be relevant to outcomes in PD^57,58^. Previous work has shown that microstructural measures, specifically fibre density correlate with white matter hyperintensities, (a key imaging signature of small vessel disease). Our findings suggest that this is relatively spared and macrostructural changes in fibre-cross section (reflecting a neurodegenerative process) are most affected^33,59^. This could be studied in future work.

Participants with PD were assessed and underwent neuroimaging on their usual dopaminergic medications to limit participant discomfort with “OFF” effects. Levodopa does not affect fractional anisotropy^60^ and is unlikely to affect fixel-based or structural connectivity metrics. However several studies have shown normalisation of functional connectivity changes in PD patients with levodopa compared to controls^61–63^. Levodopa equivalent doses did not differ between PD patients with poor vs good outcomes and dopaminergic transmission is less strongly implicated in the development of cognitive impairment in PD^64^ however further longitudinal studies of functional connectivity ON and OFF levodopa may be able to clarify whether the lack of static functional connectivity alterations reflects compensatory reorganisation or a treatment effect. Pasma biomarkers were only available from follow up sessions and not at baseline in a subset of participants; this limits the interpretation of these as early biomarkers of poor outcomes.

Assays for other phosphorylation targets of p-tau are also emerging: p-tau217 assays have recently been shown to be more sensitive to PET-amyloid positivity and subsequent progression to Alzheimer’s dementia in patients with MCI^65,66^ than p-tau181. Plasma p-tau217 levels are also predictive of abnormal tau-PET and β-amyloid CSF status in dementia with Lewy bodies and PD dementia^67^ but these have not yet been applied in earlier stages of PD.

### Conclusions

Here, we examine the changes in neuroimaging and plasma biomarkers in patients with PD who have a poorer clinical outcome at 3-year follow-up. We show extensive macrostructural white matter alterations (reduction in fibre cross-section) and reduced structural connectivity primarily in interhemispheric frontal, parietal and right occipital connectivity in those patients with poor outcomes, in the absence of gray matter or functional connectivity changes.

Increased level of plasma NFL was also found in PD with poor outcomes, correlating to white matter changes. Our study supports the further study of white matter macrostructural measures and plasma NFL as biomarkers of poor outcomes before established PD dementia occurs; and provides insight towards potential underlying processes affecting axonal tracts at early stages in the progression to PD dementia.

## Notes

### Competing Interest Statement

The authors have declared no competing interest.

