## Supplementary Material for "Neuroimaging and plasma marker evidence for white matter macrostructure loss in Parkinson’s disease"

**Figure S1. Image quality metrics per group.**

None were significantly different across the three groups.

CJV: Coefficient of joint variation, TSNR: Total signal to noise ratio, EFC: Entropy focus criterion, FD: Framewise displacement. PD: Parkinson’s disease
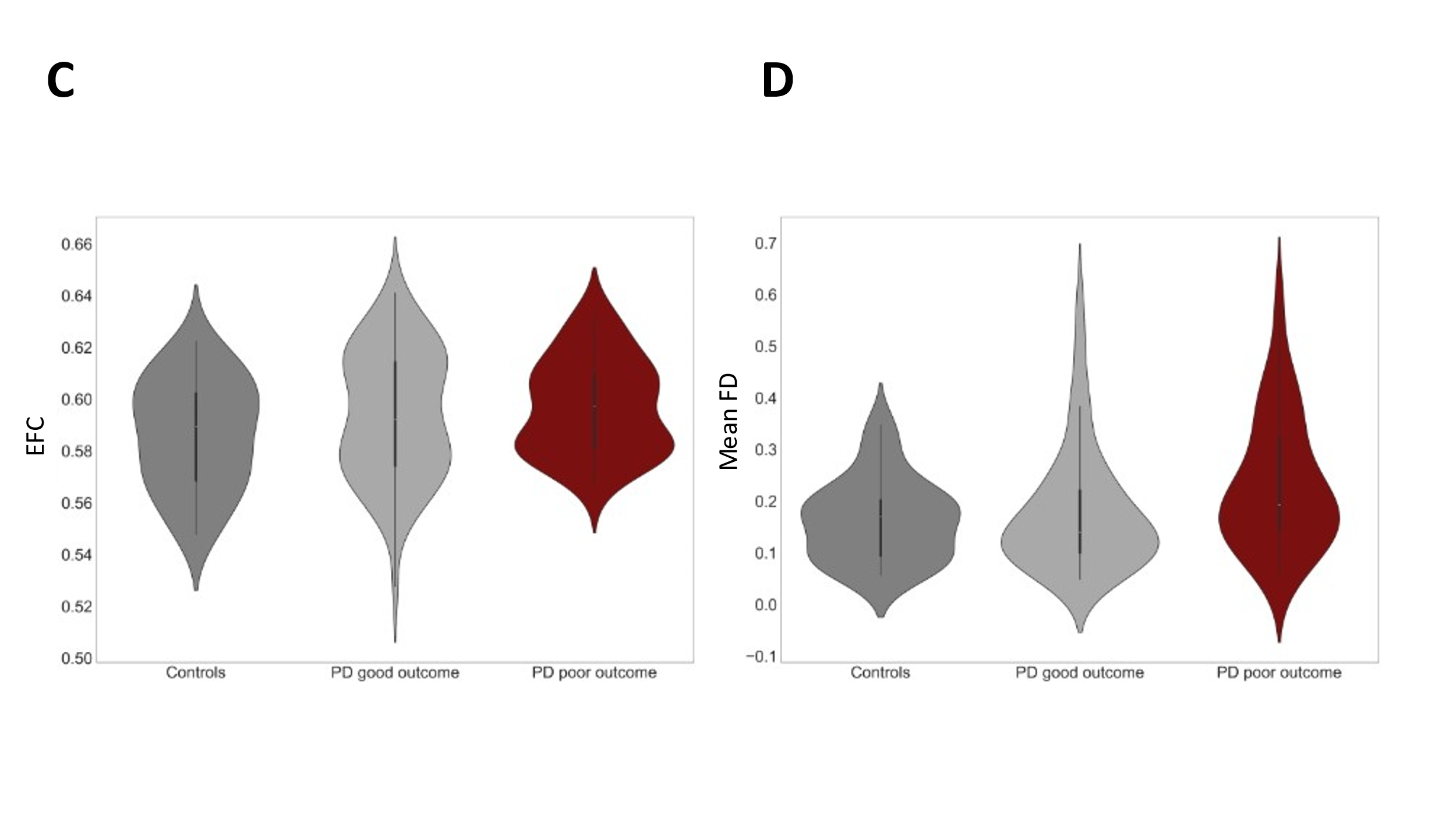

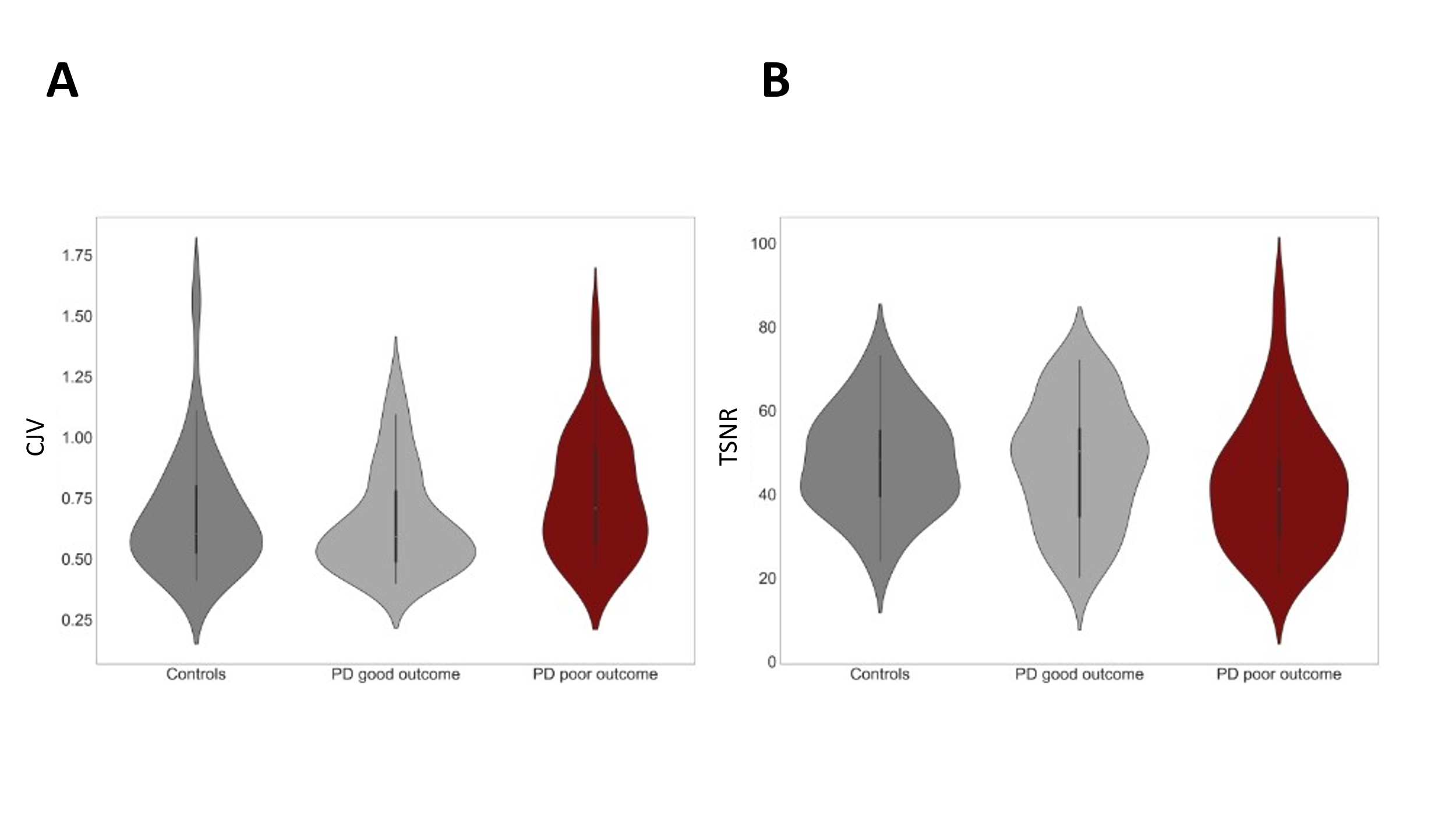

**Figure S2. Tract of interest analysis, full results.**

Mean fibre cross-section (FC) along 52 white matter tracts, segmented using TractSeg, were compared between PD with poor outcomes vs PD with poor outcomes at baseline, correcting for age, gender and total intracranial volume, false discovery rate (FDR) corrected for multiple comparisons. In colour results that survive correction for multiple comparisons (FDR-corrected p<0.05), presented as percentage change from PD with good outcomes.

AF: arcuate fasciculus, ATR: anterior thalamic radiation, CA: commissure anterior, CC: corpus callosum, CC_1: rostrum, CC_2: genu, CC_3: rostral body (Premotor), CC_4: anterior midbody (Primary Motor), CC_5: posterior midbody (Primary Somatosensory), CC_6: isthmus), CC_7: splenium, CG: cingulum, CST: corticospinal tract, MLF: middle longitudinal fascicle, FPT: fronto-pontine tract, ICP: inferior cerebellar peduncle, IFO: inferior occipito-frontal fascicle, ILF: inferior longitudinal fascicle, MCP: middle cerebellar peduncle, OR: optic radiation, POPT: parieto‐occipital pontine, SCP: superior cerebellar peduncle, SLF: superior longitudinal fascicle, STR: superior Thalamic Radiation, UF: uncinate fascicle, T_PREF: thalamo-prefrontal, T_PREM: thalamo-premotor, T_PREC: thalamo-precentral, T_POSTC: thalamo-postcentral, T_PAR: thalamo-parietal, T_OCC: thalamo-occipital.

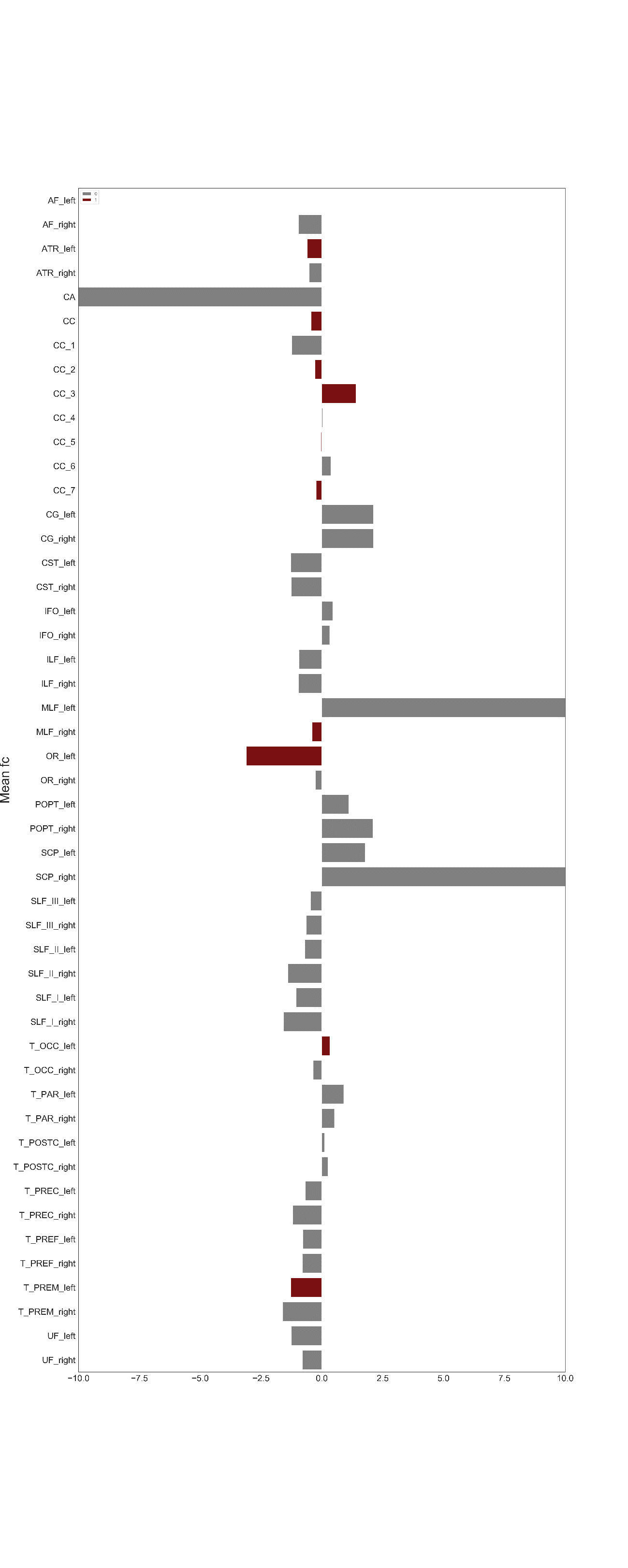

**Table S1. Network of reduced structural connectivity strength in patients with Parkinson’s disease (PD) and poor outcomes - Significant Connections, FDR-corrected p<0.05, threshold t=3.0, 5000 permutations.**

| L_V2 to L_RertroSplenialCortex. Test stat: 3.46  L_V3 to L_RertroSplenialCortex. Test stat: 3.10  L_V3A to L_PresylvianLanguage. Test stat: 3.43  L_V2 to L_ParetoOccipitalSulcus_1. Test stat: 3.09  L_RertroSplenialCortex to L_ventral_23ab. Test stat: 3.11  L_7m to L_dorsal_23ab. Test stat: 3.11  L_RertroSplenialCortex to L_8BM. Test stat: 3.04  L_RertroSplenialCortex to L_9m. Test stat: 3.55  L_10r to L_9m. Test stat: 3.12  L_8Ad to L_9m. Test stat: 3.57  L_47m to L_47l. Test stat: 3.50  L_9m to L_a9-46v. Test stat: 3.53  L_IFSa to L_11l. Test stat: 3.16  L_SuperiorTemporalVisual to L_LateralIntralParietalDorsal. Test stat: 3.08  L_9m to L_i6-8. Test stat: 3.49  L_ventral_23ab to L_PFcm. Test stat: 3.12  L_PrimaryMotorCortex_4 to L_FrontalOPercular_4. Test stat: 3.22  L_10pp to L_FrontalOPercular_4. Test stat: 3.19  L_47s to L_AnteriorAngranularInsulaComplex. Test stat: 4.23  L_11l to L_FrontalOPercular_3. Test stat: 3.35  L_AnteriorVentralInsularArea to L_FrontalOPercular_3. Test stat: 3.44  L_47m to L_AnteriorIntraParietal. Test stat: 3.01  L_V1 to L_ProStriate. Test stat: 3.26  L_LateralIntralParietalDorsal to L_ParaBeltComplex. Test stat: 3.12  L_AnteriorIntraParietal to L_ParaBeltComplex. Test stat: 3.02  L_AnteriorIntraParietal to L_AuditoryComplex_5. Test stat: 3.09  L_ProStriate to L_ParaHippocampalArea_1. Test stat: 3.04  L_ventral_23ab to L_STSd_anterior. Test stat: 3.14  L_ventral_23ab to L_STSd_posterior. Test stat: 3.22  L_LateralIntralParietalDorsal to L_TG_dorsal. Test stat: 3.08  L_V2 to L_TE2_posterior. Test stat: 3.09  L_ventral_23ab to L_TE2_posterior. Test stat: 3.28  L_V6 to L_TemporoParietoOccipitalJunction_1. Test stat: 3.08  L_V3A to L_TemporoParietoOccipitalJunction_1. Test stat: 3.78  L_PFcm to L_DorsalTransitionalVisualArea. Test stat: 3.29  L_FrontalOPercular_1 to L_IntraParietal_1. Test stat: 3.29  L_TE2_posterior to L_IntraParietal_1. Test stat: 3.37  L_DorsalTransitionalVisualArea to L_IntraParietal_1. Test stat: 3.32  L_V3A to L_PF. Test stat: 3.02  L_TE2_posterior to L_PF. Test stat: 3.65  L_IntraParietal_1 to L_V6A. Test stat: 3.08  L_V1 to L_FST. Test stat: 3.14  L_44 to L_LateralOccipital_3. Test stat: 3.00  L_AnteriorAngranularInsulaComplex to L_FrontalOpercular_5. Test stat: 3.03  L_9m to L_p47r. Test stat: 3.64  L_LateralIntralParietalDorsal to L_AuditoryComplex_4. Test stat: 3.13  L_7PC to L_STSv_anterior. Test stat: 3.27  L_LateralIntralParietalDorsal to L_STSv_anterior. Test stat: 3.27  L_V1 to R_V1. Test stat: 3.38  L_ProStriate to R_V6. Test stat: 3.30  L_23c to R_PrimaryMotorCortex_4. Test stat: 3.62  L_24d_dorsal to R_PrimaryMotorCortex_4. Test stat: 3.28  L_24d_ventral to R_PrimaryMotorCortex_4. Test stat: 3.21  L_LateralIntralParietalDorsal to R_ParietoOccipitalSulcus_2. Test stat: 3.41  L_7P_lateral to R_PrimaryAuditory_1. Test stat: 3.36  L_7m to R_PrecuneusVisual. Test stat: 3.06  L_5m to R_PrecuneusVisual. Test stat: 3.40  L_6mp to R_PrecuneusVisual. Test stat: 3.40  L_ProStriate to R_7Pm. Test stat: 3.03  L_5m to R_7m. Test stat: 3.01  L_5L to R_7m. Test stat: 3.27  L_7A_medial to R_7m. Test stat: 3.17  L_5L to R_ParetoOccipitalSulcus_1. Test stat: 3.25  L_7PC to R_ParetoOccipitalSulcus_1. Test stat: 3.47  L_VentralIntraParietalComplex to R_ParetoOccipitalSulcus_1. Test stat: 3.22  L_PF to R_ParetoOccipitalSulcus_1. Test stat: 3.20  L_5L to R_ventraR_23ab. Test stat: 3.24  L_PrimaryMotorCortex_4 to R_dorsaR_23ab. Test stat: 3.60  L_55b to R_dorsaR_23ab. Test stat: 3.21  L_2 to R_dorsaR_23ab. Test stat: 3.11  L_6mp to R_dorsaR_23ab. Test stat: 3.02  L_PrimarySenroryCortex_3b to R_31p_ventral. Test stat: 3.12  L_FrontalEyeFields to R_31p_ventral. Test stat: 3.24  L_7A_medial to R_31p_ventral. Test stat: 3.32  L_2 to R_31p_ventral. Test stat: 3.00  L_6mp to R_31p_ventral. Test stat: 3.05  L_PFt to R_31p_ventral. Test stat: 3.12  L_ParaBeltComplex to R_31p_ventral. Test stat: 3.10  L_24d_dorsal to R_5m. Test stat: 3.72  L_SuperiorFrontalLanguage to R_24d_dorsal. Test stat: 3.29  L_23c to R_24d_dorsal. Test stat: 3.22  L_24d_ventral to R_24d_dorsal. Test stat: 4.42  L_SupplementaryCingulateEyeField to R_24d_dorsal. Test stat: 3.22  L_6ma to R_24d_dorsal. Test stat: 3.21  L_p32pr to R_24d_dorsal. Test stat: 3.15  L_6a to R_24d_dorsal. Test stat: 3.41  L_PrimaryMotorCortex_4 to R_7_lateraR_area. Test stat: 3.01  L_V3A to R_7_lateraR_area. Test stat: 3.16  L_ventral_23ab to R_7_lateraR_area. Test stat: 3.14  L_5m to R_7_lateraR_area. Test stat: 3.43  L_6r to R_7_lateraR_area. Test stat: 3.13  L_24d_ventral to R_SupplementaryCingulateEyeField. Test stat: 3.13  L_11l to R_SupplementaryCingulateEyeField. Test stat: 3.58  L_24d_ventral to R_6ma. Test stat: 3.15  L_V1 to R_7A_medial. Test stat: 3.50  L_V2 to R_7A_medial. Test stat: 3.59  L_RertroSplenialCortex to R_7A_medial. Test stat: 3.02  L_7m to R_7A_medial. Test stat: 3.42  L_ventral_23ab to R_7A_medial. Test stat: 3.00  L_24d_ventral to R_7A_medial. Test stat: 3.20  L_7P_lateral to R_7A_medial. Test stat: 3.06  L_d32 to R_7A_medial. Test stat: 3.53  L_44 to R_7A_medial. Test stat: 3.10  L_24d_ventral to R_7P_lateral. Test stat: 3.37  L_V3 to R_7PC. Test stat: 3.02  L_V1 to R_MedialIntralParietal. Test stat: 4.05  L_V2 to R_MedialIntralParietal. Test stat: 3.49  L_ventral_23ab to R_2. Test stat: 3.22  L_LateralOccipital_3 to R_2. Test stat: 3.21  L_a32pr to R_2. Test stat: 3.28  L_SuperiorFrontalLanguage to R_6mp. Test stat: 3.14  L_24d_ventral to R_6mp. Test stat: 3.72  L_SupplementaryCingulateEyeField to R_6mp. Test stat: 3.25  L_8Ad to R_p32pr. Test stat: 3.10 | L_Caudate to L_Putamen. Test stat: 3.52  L_47s to L_Pallidum. Test stat: 3.20  L_Caudate to L_Pallidum. Test stat: 3.19  L_FrontalOpercular_5 to Brain-Stem. Test stat: 3.22  L_p47r to Brain-Stem. Test stat: 3.10  R_43 to Brain-Stem. Test stat: 3.09  L_V3 to L_Hippocampus. Test stat: 3.02  L_V8 to L_Hippocampus. Test stat: 3.26  L_SuperiorTemporalVisual to L_Hippocampus. Test stat: 3.10  L_7A_medial to L_Hippocampus. Test stat: 3.37  L_7PC to L_Hippocampus. Test stat: 3.23  L_8Av to L_Hippocampus. Test stat: 3.60  L_8C to L_Hippocampus. Test stat: 3.00  L_44 to L_Hippocampus. Test stat: 3.00  L_6r to L_Hippocampus. Test stat: 3.01  L_s6-8 to L_Hippocampus. Test stat: 3.01  L_OP4 to L_Hippocampus. Test stat: 3.14  L_TG_dorsal to L_Hippocampus. Test stat: 3.32  L_PF to L_Hippocampus. Test stat: 3.65  L_FST to L_Hippocampus. Test stat: 3.80  L_p47r to L_Hippocampus. Test stat: 3.00  L_TG_ventral to L_Hippocampus. Test stat: 3.56  R_PrimaryAuditory_1 to L_Hippocampus. Test stat: 3.09  R_7A_medial to L_Hippocampus. Test stat: 3.12  L_Thalamus to L_Hippocampus. Test stat: 3.39  L_ventral_23ab to L_Amygdala. Test stat: 3.02  R_LateralIntraParietalVentral to L_Amygdala. Test stat: 3.03  L_Thalamus to L_VentralDC. Test stat: 3.58  R_8C to R_Thalamus. Test stat: 3.11  R_s6-8 to R_Thalamus. Test stat: 3.45  R_43 to R_Thalamus. Test stat: 3.19  R_FrontalOPercular_4 to R_Thalamus. Test stat: 3.25  R_8Av to R_Caudate. Test stat: 3.25  R_8Ad to R_Caudate. Test stat: 3.21  R_45 to R_Caudate. Test stat: 3.34  R_47l to R_Caudate. Test stat: 3.71  R_a47r to R_Caudate. Test stat: 3.48  R_a9-46v to R_Caudate. Test stat: 3.37  R_9-46d to R_Caudate. Test stat: 3.27  R_AnteriorIntraParietal to R_Caudate. Test stat: 3.21  R_IntraParietaR_2 to R_Caudate. Test stat: 3.19  R_IntraParietaR_1 to R_Caudate. Test stat: 3.71  R_FST to R_Caudate. Test stat: 3.22  R_p47r to R_Caudate. Test stat: 3.97  R_p47r to R_Putamen. Test stat: 3.01  Brain-Stem to R_Putamen. Test stat: 3.49  R_Thalamus to R_Putamen. Test stat: 3.15  R_Caudate to R_Putamen. Test stat: 3.21  R_43 to R_Pallidum. Test stat: 3.36  L_7A_medial to R_Hippocampus. Test stat: 3.03  L_7PC to R_Hippocampus. Test stat: 3.40  R_TG_dorsal to R_Hippocampus. Test stat: 3.83  R_TE1_middle to R_Hippocampus. Test stat: 3.03  R_Thalamus to R_Hippocampus. Test stat: 3.27  R_PF to R_Amygdala. Test stat: 3.55  R_Caudate to R_Amygdala. Test stat: 3.37  R_SupplementaryCingulateEyeField to R_VentralDC. Test stat: 3.04  R_43 to R_VentralDC. Test stat: 3.57  R_OP4 to R_VentralDC. Test stat: 3.17  R_IntraParietaR_2 to R_VentralDC. Test stat: 3.16  R_PF_opercular to R_VentralDC. Test stat: 3.20  R_LateralIntralParietalDorsal to R_43. Test stat: 3.07  R_9-46d to R_OP2-3. Test stat: 3.04  L_10r to R_MiddleInsular. Test stat: 3.01  R_LateralIntralParietalDorsal to R_FrontalOPercular_2. Test stat: 3.03  L_p32pr to R_AnteriorIntraParietal. Test stat: 3.28  R_PFt to R_AnteriorIntraParietal. Test stat: 3.24  L_VentralIntraParietalComplex to R_PreSubiculum. Test stat: 3.07  R_7PC to R_PreSubiculum. Test stat: 3.05  L_5m to R_ParaBeltComplex. Test stat: 3.35  L_V1 to R_ParaHippocampalArea_1. Test stat: 3.30  L_ProStriate to R_ParaHippocampalArea_1. Test stat: 3.19  R_7A_medial to R_TE1_posterior. Test stat: 3.04  L_LateralIntralParietalDorsal to R_TemporoParietoOccipitalJunction_1. Test stat: 3.01  R_43 to R_IntraParietaR_1. Test stat: 3.25  L_V2 to R_IntraParietaR_0. Test stat: 3.33  R_SuperiorTemporalVisual to R_IntraParietaR_0. Test stat: 3.60  R_AnteriorIntraParietal to R_PF_opercular. Test stat: 3.42  L_5m to R_31pd. Test stat: 3.01  L_7_lateral_area to R_31pd. Test stat: 3.11  L_8Ad to R_posteriorOFC. Test stat: 3.74  L_p47r to R_posteriorOFC. Test stat: 3.31  R_AnteriorAngranularInsulaComplex to R_p47r. Test stat: 3.16  R_V3A to R_LateralBeltComplex. Test stat: 3.08  R_9-46d to R_LateralBeltComplex. Test stat: 3.07  R_InsularGranularComplex to R_LateralBeltComplex. Test stat: 3.12  R_LateralIntralParietalDorsal to R_TE1_middle. Test stat: 3.03  L_8BL to R_a32pr. Test stat: 3.01  L_a10p to R_a32pr. Test stat: 3.23  L_p47r to L_Cerebellum. Test stat: 3.01  L_SuperiorFrontalLanguage to L_Caudate. Test stat: 3.22  L_SuperiorTemporalVisual to L_Caudate. Test stat: 3.12  L_SupplementaryCingulateEyeField to L_Caudate. Test stat: 3.41  L_6ma to L_Caudate. Test stat: 3.09  L_8Av to L_Caudate. Test stat: 3.62  L_8C to L_Caudate. Test stat: 3.30  L_47l to L_Caudate. Test stat: 3.28  L_47s to L_Caudate. Test stat: 3.46  L_i6-8 to L_Caudate. Test stat: 3.27  L_TE1_posterior to L_Caudate. Test stat: 3.09  L_DorsalTransitionalVisualArea to L_Caudate. Test stat: 3.10  L_p47r to L_Caudate. Test stat: 3.23  L_AuditoryComplex_4 to L_Caudate. Test stat: 3.05  L_47s to L_Putamen. Test stat: 4.14  L_FrontalOpercular_5 to L_Putamen. Test stat: 3.04  R_SupplementaryCingulateEyeField to R_p32pr. Test stat: 3.24  L_a9-46v to R_a24. Test stat: 3.21  L_a9-46v to R_10r. Test stat: 3.25  L_a9-46v to R_9m. Test stat: 3.23  R_5L to R_a47r. Test stat: 3.14  L_VentralIntraParietalComplex to R_IFJp. Test stat: 3.32  L_a32pr to R_9-46d. Test stat: 3.10  L_8Ad to R_a10p. Test stat: 3.67  L_p47r to R_a10p. Test stat: 3.33  L_a32pr to R_10pp. Test stat: 3.28  L_8Ad to R_OrbitoFrontalCortex. Test stat: 3.11  L_7P_lateral to R_LateralIntralParietalDorsal. Test stat: 3.53  R_6r to R_LateralIntralParietalDorsal. Test stat: 3.44  L_p32pr to R_s6-8. Test stat: 3.07  R_a10p to R_s6-8. Test stat: 3.38  L_8BL to R_p32pr. Test stat: 3.18 |
| --- | --- |
